# LSTrAP-Crowd: Prediction of novel components of bacterial ribosomes with crowd-sourced analysis of RNA sequencing data

**DOI:** 10.1101/2020.04.20.005249

**Authors:** Benedict Hew, Qiao Wen Tan, William Goh, Jonathan Wei Xiong Ng, Kenny Koh, Ryan Chieh Feng Rugdee, Zheng Kai Teng, Jun Xiong Tan, Xi Yei, Qing Rong Tan, Ifa Syafiqah Binte Sulaiman, Seo Min Li Gilia, Erielle Marie Fajardo Villanueva, Son Thanh Nguyen, Dhira Anindya Putri, Jovi Tan Siying, Teo Yong Ren Johanan, Jia Wei Tan, Koh Shao Ning, Gladys, Wei Wen Ong, Jia Rong Moo, Jace Koh, Pei Xuan Lim, Shook Wei Teoh, Pravin Tamilselvam, Harvard Hui, Yi Xuan Chua, Yook Kit Ow Yeong, Tay Jian Hua, Ming Jun Chong, Yu Wei Sho, Bridget Jing Xing Tang, Carissa Yuwono Kwantalalu, Nur Afiqah Binte Mohammad Rizal, Wei Heng Tan, Lim Shan Chun, Sherianne Yen Tze Tan, Tan Jia Ying, Audrey Michelle Luminary, Lim Jia Jia, Jolyn, Vanessa Lunardi, Ann Don Low, M K Abdul Rahim, Lin Ming, Joseph JQ Ng, Han Tsou, Cheryl Lim Jiayi, Teffarina Tay Hui Wen, Valerie Teo Fang Wei, Tan You Sheng Justin, Shellia Oktavina, Aaminatul Khalishah Binte Roslan, Natasha Cassandra Chee, Zoe Chen Hui Xin, Nhi Uyen Le Nguyen, Tristy Abigayle Marta, Poh Jia’En, Ang Wei Ying, Alena Tay Qi Ye, Chiu Wei Yeow Emile, Wong Xanaz, Xylon Wei Rui Lee, Wong Pei Wen Kelly, Zhe Jun Tan, Vishalini Val R, Rayna Yip, Cherlyn Chua, Kai Lun Boon, Sriya Mulupuri, Lim Yuen Shan, Samantha Chee Suhui, Amanda Crystal Lee Wei Jin, Siew Choo Tey, Qi Ying Neo, Chan Yi Hue, Phua Tian Xin, Ana Ho Sze Qi, Edbert E. Rodrigues, Chan Mu En, Dynn Sim, Marcus Chee, Abigail Ho, Ang Wen hui, Bertrand Wong, Margaret X Zhao, Er Kian Ching Gabbie, Deng Zheyun Grace, Xin Yi Tan, Melissa Foong, Lim Qin Rui Rachel, Alyssa Jiaqi Lim, Seow Jia Xuan, Rinta P. Reji, Devika Menon, Ong Xuan Xuan, Nicole, Ravi Keerthana, Min Jia Wong, Zachary J D’Rozario, Shing Yee Lim, Nicholas Lee, Ying Ni, Ying Lian, Jing Wen Poh, Ming Jern Adrian Lee, Pravenah Ravi Chandran, Jia Xin Ngiaw, Herman Foo, Joash Poon, Tong Ling Chan, Perry Ng, Ashlyn Xuanqi Ng, Zhen Ying Ong, Lee Wan Xuan Trena, Lim Shi Min Kristy, Yu Xuan Thng, Ong Si Yi Shirley, Sau Thi Chu, Shu Hua Samantha Lim, Jun Sheng Ho, Celest Lixuan Phang, Victoria Toh Le Yi, Peiran Ng, Seetoh Wei Song, Manessa Nah Shue Ern, Marek Mutwil

## Abstract

Bacterial resistance to antibiotics is a growing problem that is projected to cause more deaths than cancer in 2050. Consequently, novel antibiotics are urgently needed. Since more than half of the available antibiotics target the bacterial ribosomes, proteins that are involved in protein synthesis are thus prime targets for the development of novel antibiotics. However, experimental identification of these potential antibiotic target proteins can be labor-intensive and challenging, as these proteins are likely to be poorly characterized and specific to few bacteria. In order to identify these novel proteins, we established a Large-Scale Transcriptomic Analysis Pipeline in Crowd (LSTrAP-Crowd), where 285 individuals processed 26 terabytes of RNA-sequencing data of the 17 most notorious bacterial pathogens. In total, the crowd processed 26,269 RNA-seq experiments and used the data to construct gene co-expression networks, which were used to identify more than a hundred uncharacterized genes that were transcriptionally associated with protein synthesis. We provide the identity of these genes together with the processed gene expression data. The data can be used to identify other vulnerabilities or bacteria, while our approach demonstrates how the processing of gene expression data can be easily crowdsourced.

## Introduction

Bacterial resistance to antibiotics is a serious and growing concern in public health, taking ca. 99,000 lives and costing 21-34 billion USD per year in the USA (Spellberg *et al*., 2011). Methicillin-resistant Gram-positive *Staphylococcus aureus* (MRSA) and Gram-negative *Pseudomonas aeruginosa* are the leading causes of serious infections as they form biofilms. Biofilms are complex bacterial communities embedded in an extracellular matrix, and these communities are able to resist antimicrobial agents (Mah and O’Toole, 2001). For instance, bacteria can be up to 1000x more tolerant to antibiotics when they grow as a biofilm, compared to single cell suspension (planktonic cells). Consequently, new antibiotics are urgently needed to combat these resistance mechanisms, either alone or in combination with existing drugs.

More than half of the antibiotics currently in use target the bacterial ribosome, typically at the elongation step of protein synthesis (Wilson, 2014), through direct or proximal binding of the peptidyl transferase center (PTC) which catalyzes peptide bond formation (Arenz and Wilson, 2016). PTCL□targeting antibiotics (e.g., lincosamides, pleuromutilins, chloramphenicol and group A streptogramins), inhibit protein synthesis by obstructing the proper positioning of the tRNA substrates (Dunkle *et al*., 2010).

Bacteria can be intrinsically less sensitive to antibiotics due to less efficient uptake of antibiotics or mutations in ribosomal proteins that result in decreased drug□binding efficiency (Wilson, 2014; Dinos, 2017). The most frequently encountered acquired resistance mechanism involves the methylation of the ribosomal RNA (e.g., by Erm family methyltransferases), which results in decreased drug□binding efficiency and increased viability in the presence of antibiotics (Wilson, 2009; Liu and Douthwaite, 2002). As modification of the ribosomes can result in a decrease in fitness, these methyltransferases genes tend to be induced by the relevant antibiotics through translation attenuation (Lin *et al*., 2018; Vazquez-Laslop *et al*., 2008). Alternatively, the antibiotics can also be modified, pumped out, or degraded, thus lowering the intracellular concentration to non□toxic levels (Wilson, 2014; Golkar *et al*., 2018). Another mechanism is ribosome protection, where the antibiotic is actively dislodged from the ribosome by ATP□binding cassette F (ABC□F) protein, as observed in many clinical isolates (e.g., *Pseudomonas aeruginosa, Escherichia coli, Staphylococcus aureus, Enterococcus faecalis* and *Listeria monocytogenes*)(Sharkey and O’Neill, 2018; Murina *et al*., 2019; Kerr *et al*., 2005).

While the structure of the ribosomes is well conserved, structural features of ribosomes may vary significantly between different species, suggesting species-specific adaptations of protein synthesis (Ahmed *et al*., 2016; Ahmed *et al*., 2017; Barandun *et al*., 2019; Bieri *et al*., 2017; Kushwaha and Bhushan, 2020; Eyal *et al*., 2015; Greber and Ban, 2016; Melnikov *et al*., 2012; Melnikov *et al*., 2018). For example, structural analysis of mycobacterial ribosome revealed that the 30S ribosomal subunit lacks the protein bS21 that is found in *Escherichia coli*. Instead, the mycobacteria employ a unique protein bS22 near the decoding center (DC), thereby keeping the overall number of ribosomal proteins in 30S subunit the same as in *E. coli* (Kushwaha and Bhushan, 2020). Thus, the identification of novel bacteria ribosomal components has great potential for the development of species-specific antibiotics. However, the identification of these novel components using traditional molecular or structural biology approaches is time-consuming.

Bioinformatic approaches are used to predict gene function and can be used to identify novel components of protein synthesis. Newly sequenced genomes of all organisms are typically first annotated using sequence similarity analysis, where the genes are annotated based on the DNA/protein sequence similarity to characterized genes/proteins (Rhee and Mutwil, 2014). While sequence similarity analysis is well established and gives a quick overview of gene functions in a new genome, it has its caveats as genes can i) have multiple functions, ii) sub- or neo-functionalise and/or iii) have no sequence similarity to characterized genes. Thus, while sequence similarity analysis is a powerful method, it requires other methods to complement it (Rhee and Mutwil, 2014; Proost and Mutwil, 2016).

The wide availability of RNA sequencing (RNA-seq) data makes it possible to study gene function from the perspective of gene expression (Rhee and Mutwil, 2014; Usadel *et al*., 2009; Hansen *et al*., 2018; Hansen *et al*., 2014). Co-expression analysis is based on the observation that genes that have similar expression profiles across experiments tend to be functionally related (Rhee and Mutwil, 2014; Proost and Mutwil, 2016; Proost and Mutwil, 2017). These co-expressed genes can be identified by analyzing publicly-available microarrays or RNA-seq data, and the co-expression relationships can be represented as networks. In a co-expression network, genes are represented as nodes, where edges connect co-expressed nodes (links)(Mutwil *et al*., 2008; Mutwil *et al*., 2009; Mutwil *et al*., 2010; Mutwil *et al*., 2011; Ruprecht *et al*., 2016; Ferrari *et al*., 2018; Ruprecht *et al*., 2017; Ng *et al*., 2019; Wen Tan and Mutwil, 2019; Proost and Mutwil, 2018a; Ruprecht *et al*., 2011; Proost and Mutwil, 2018b). The networks can be mined for groups of highly connected genes (called clusters or modules) that likely represent genes that are involved in the same biological process. Due to the ubiquity of expression data, and the ability to complement DNA/protein sequence-based gene function prediction approaches, coexpression networks have become a popular tool to elucidate the function of genes. The networks have predicted the function of genes involved in a wide range of processes, such as various cellular processes (Takabayashi *et al*., 2009; Takahashi *et al*., 2008; Stuart *et al*., 2003; Wen Tan and Mutwil, 2019), transcriptional regulation (Yu *et al*., 2003), physiological responses to the environment and stress (Lee *et al*., 2010; Jiménez-Gómez *et al*., 2010), and the biosynthesis of metabolites (Tan *et al*., 2020; Tan and Mutwil, 2019; Ruprecht *et al*., 2016; Sibout *et al*., 2017).

The amount of gene expression data has expanded vastly over the last decade, resulting in >1000-fold increase in nucleotide bases on NCBI Sequence Read Archive (SRA), from 11TB (2010) to 12 PB (2020). Due to limitations in software used to estimate gene expression from RNA-seq data, analyzing all this data would have been unthinkable a decade ago. However, drastic improvements to the speed and efficiency of software, such as Kallisto (Bray *et al*., 2016) and salmon (Patro *et al*., 2017), allow the analysis of gigabytes of data on even a Raspberry Pi-like miniature computer (Tan and Mutwil, 2019). Recently, combined the availability of cloud computing and the user-friendliness of the Jupyter notebooks to implement a large-scale transcriptomic analysis pipeline, LSTrAP-Cloud (Tan *et al*., 2020). Importantly, though Google Colab, the pipeline gives access to a free cloud computer with 2 Xeon cores, with at least 15 gigabytes of permanent storage (as provided by users google drive account) and 12 gigabytes of RAM, giving biologists both the software and hardware to perform large-scale co-expression analysis.

In this study, we introduce Large-Scale Transcriptomic Analysis Pipeline in Crowd (LSTrAP-Crowd). This simple pipeline was used by 285 undergraduate students to process RNA-seq data of some of the 17 most notorious bacterial pathogens. Within a week, the students processed 26,269 RNA-seq samples, comprising 263,757,103,900 (∼263 billion) reads and 26.38 terabytes of data. The gene expression data was used to construct co-expression networks, which were mined for the presence of uncharacterized genes that were co-expressed with the bacterial ribosomes. In total, we have predicted more than 100 putative proteins to be involved in protein synthesis in the 17 bacterial pathogens.

## Materials and methods

### Streaming RNA Sequencing Data

The LSTrAP-Crowd pipeline was implemented on Google Colaboratory and is based on the LSTrAP-Cloud pipeline with standard parameters (Tan *et al*., 2020). The pipeline streams the RNA-seq fastq files to a virtual machine in the cloud, and deposits the processed gene expression data on the user’s Google Drive. The CDSs were obtained from EnsembleGenomes and used to generate kallisto index with default parameters. The RNA sequencing data of the 17 bacteria was obtained from European Nucleotide Archive (ENA) and mapped against the kallisto index of coding sequences (CDS) of the 17 bacteria. The used CDSs are: *Campylobacter jejuni* (Campylobacter_jejuni_subsp_jejuni_cg8421.ASM17179v2.cds.all.fa), *Clostridioides difficile* (Clostridioides_difficile_e25.E25.cds.all.fa), *Enterococcus faecalis* (Enterococcus_faecalis_og1rf.ASM17257v2.cds.all.fa), *Escherichia coli* (Escherichia_coli_str_k_12_substr_mg1655.ASM584v2.cds.all.fa), *Haemophilus influenzae* (Haemophilus_influenzae_r3021.ASM16975v1.cds.all.fa), *Helicobacter pylori* (Helicobacter_pylori_b8.ASM19675v1.cds.all.fa), *Klebsiella pneumoniae* (Klebsiella_pneumoniae_jm45.ASM44540v1.cds.all.fa), *Listeria monocytogenes* (Listeria_monocytogenes_gca_001027125.ASM102712v1.cds.all.fa), *Mycobacterium tuberculosis* (Mycobacterium_tuberculosis_h37rv.ASM19595v2.cds.all.fa), *Mycoplasma pneumoniae* (Mycoplasma_pneumoniae_fh.ASM14394v1.cds.all.fa), *Neisseria gonorrhoeae* (Neisseria_gonorrhoeae_gca_001047275.ASM104727v1.cds.all.fa), *Pseudomonas aeruginosa* (Pseudomonas_aeruginosa_gca_001181725.E11_London_26_VIM_2_06_13.cds.all.fa), *Salmonella enterica* (Salmonella_enterica_subsp_enterica_serovar_typhimurium_str_lt2.ASM694v2.cds.all.f a), *Staphylococcus aureus* (Staphylococcus_aureus_gca_001212685.7738_4_69.cds.all.fa), *Streptococcus pneumoniae* (Streptococcus_pneumoniae_r6.ASM704v1.cds.all.fa), *Streptococcus pyogenes* (Streptococcus_pyogenes_ns88_2.SPNS88.2.cds.all.fa), *Vibrio cholerae* (Vibrio_cholerae_v51.ASM15246v2.cds.all.fa). A total of 26,269 experiments were streamed (Table S1).

### Generating gene expression matrices for the 17 bacteria

To remove RNA-seq samples that are of lower quality, we identified outlier samples that show a lower number (n_pseudoaligned) and percentage (p_pseudoaligned) of reads aligned to the coding sequences than the majority of the samples. This analysis assumes that the majority of samples are of good quality. The expression matrices containing the gene expression data that passed these thresholds are available in Supplementary Dataset 1. Table S1 contains the n_pseudoaligned and p_pseudoaligned numbers and indicates which samples passed the thresholds.

### Identification of genes Involved in ribosome biogenesis with co-expression networks

To identify genes that are involved in protein synthesis in the 17 bacteria, we have first retrieved all genes containing ‘DUF’ (domain of unknown function), ‘hypothetical’ or ‘conserved’ in their description. Next, we calculated the Pearson Correlation Coefficient (PCC) between the uncharacterized genes and all genes in the genome, where PCC>0.7 between two genes was used to indicate that the genes are co-expressed. Finally, the uncharacterized genes were predicted to be involved in protein synthesis if >10%, >30%, >50%, >70% or >90% of the genes that they are co-expressed with contained annotations such as ‘ribosome’ or ‘ribosomal’.

## Results

### Obtaining and quality-controlling gene expression data for 17 bacterial pathogens

In this study, we analyzed the gene expression data of 17 notorious bacterial pathogens that cause numerous diseases, such as pharyngitis, tonsillitis, scarlet fever, cellulitis, erysipelas, rheumatic fever, post-streptococcal glomerulonephritis, necrotizing fasciitis, and many others (Table 1). While more bacterial pathogens were considered, we only analyzed bacteria that had at least 100 RNA-seq samples based on Illumina technology found in the Sequence Read Archive (Leinonen *et al*., 2011). In total, 26,269 RNA-seq samples were analyzed.

**Table 1.** Genomic properties of the 17 bacteria and the RNA-seq sample statistics.

| **Bacteria** | **Number of genes** | **Genome size (Mb)** | **Student groups analyzing the data** | **RNA-seq samples: total/passed QC** |
| --- | --- | --- | --- | --- |
| *Campylobacter jejuni* | 1635 | 1.65 | 1 | 219/320 |
| *Clostridioides difficile* | 3769 | 4.27 | 1 | 182/381 |
| *Enterococcus faecalis* | 2579 | 3.11 | 1 | 81/195 |
| *Escherichia coli* | 4141 | 5.44 | 13 | 2494/7154 |
| *Haemophilus influenzae* | 2098 | 1.79 | 2 | 189/448 |
| *Helicobacter pylori* | 1720 | 1.62 | 1 | 84/167 |
| *Klebsiella pneumoniae* | 5141 | 5.78 | 2 | 536/639 |
| *Listeria monocytogenes* | 2817 | 2.78 | 2 | 370/572 |
| *Mycobacterium tuberculosis* | 4023 | 4.33 | 11 | 2414/6495 |
| *Mycoplasma pneumoniae* | 629 | 0.82 | 3 | 365/956 |
| *Neisseria gonorrhoeae* | 2159 | 2.15 | 1 | 102/371 |
| *Pseudomonas aeruginosa* | 6512 | 6.09 | 7 | 1372/2662 |
| *Salmonella enterica* | 4554 | 4.79 | 3 | 611/1284 |
| *Staphylococcus aureus* | 2638 | 2.90 | 5 | 1112/1963 |
| *Streptococcus pneumoniae* | 2043 | 2.13 | 2 | 365/956 |
| *Streptococcus pyogenes* | 1660 | 1.85 | 4 | 1340/1705 |
| *Vibrio cholerae* | 3648 | 4.00 | 1 | 167/311 |

The RNA-seq data was streamed by using a modified LSTRaP-Cloud pipeline (Figure 1A), which gives each user a free Google Colab notebook equipped with a 2 core Xeon CPU and 12 Gb of RAM (Tan *et al*., 2020). The modified pipeline, LSTrAP- Crowd, thus allows a large group of people to download the gene expression data collaboratively. 285 first-year undergraduate students were divided into 60 groups, with each group tasked to download a maximum of 600 RNA-seq samples (Table 1). The size of each RNA-seq sample was capped at ∼1 Gb, allowing a person running the modified LSTRaP-Crowd pipeline to download ∼300 RNA-seq samples per day (Tan *et al*., 2020). Theoretically, 85,500 (300*285) RNA-seq samples equivalent to ∼85Tb, could be processed per day by the classroom.

**Figure 1.**
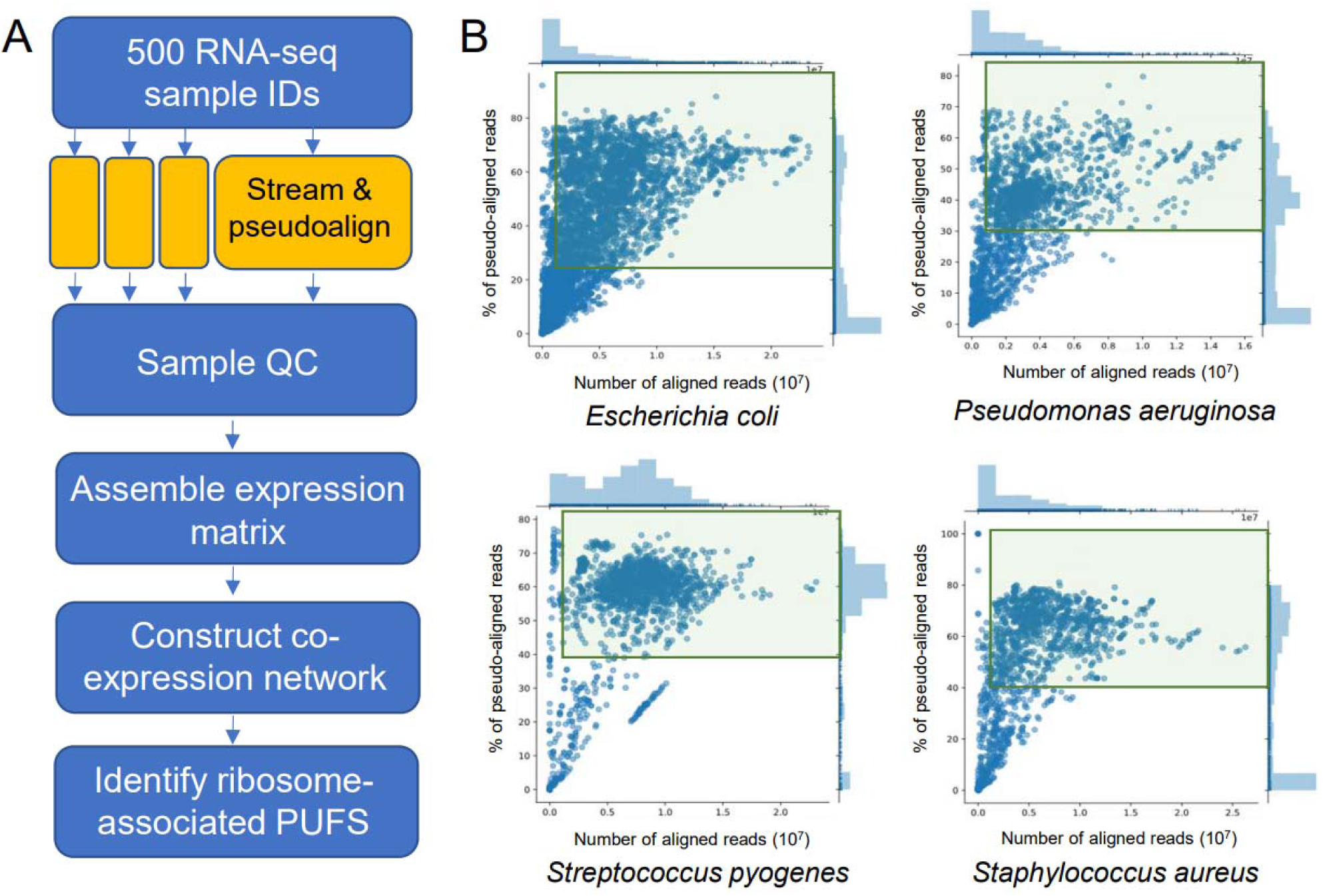
LSTrAP-Crowd pipeline and sample quality control. A) LSTrAP-Crowd pipeline. The pipeline is modification of LSTrAP-Cloud, where the modification allows one group of students to share the task streaming and pseudo-aligning the RNA-seq data. B) Scatter plot showing the number of pseudoaligned reads (n_pseudoaligned, x- axis) and % of pseudoaligned reads (p_pseudoaligned, y-axis) of RNA-seq samples for four bacteria. Samples wit n_pseudoaligned>1,000,000 and with high p_pseudoaligned values that were not dissimilar from the majority of samples were used to build the expression matrices. Used samples are indicated by green rectangles.

For each species, all the processed RNA-seq experiments were visualized as scatter plots, that show the percentage (y-axis) against the number (x-label) of read pseudoaligned to the respective species’ CDS. For each experiment, hig pseudoalignment percentage indicates high sequence similarity to the CDS, whereas high absolute number of reads indicates whether the experiment has sufficient data for meaningful coexpression analysis. In this study, a minimum threshold of 1 million read pseudoaligned was required for the experiment to be considered. We removed samples with n_pseudoaligned<1,000,000 and with p_pseudoaligned values that were lower than the majority of the high p_pseudoaligned samples (typically >30%) (Figure 2, Figure S1). The scatterplot pattern was different for each bacteria, most likely due to each bacteria having a different ratio of coding to non-coding DNA (Figure S1). Samples that passed these thresholds were used to build expression matrices (Supplementary Data 1) and used for the co-expression analysis and identification of novel genes involved in protein synthesis.

**Figure 2.**
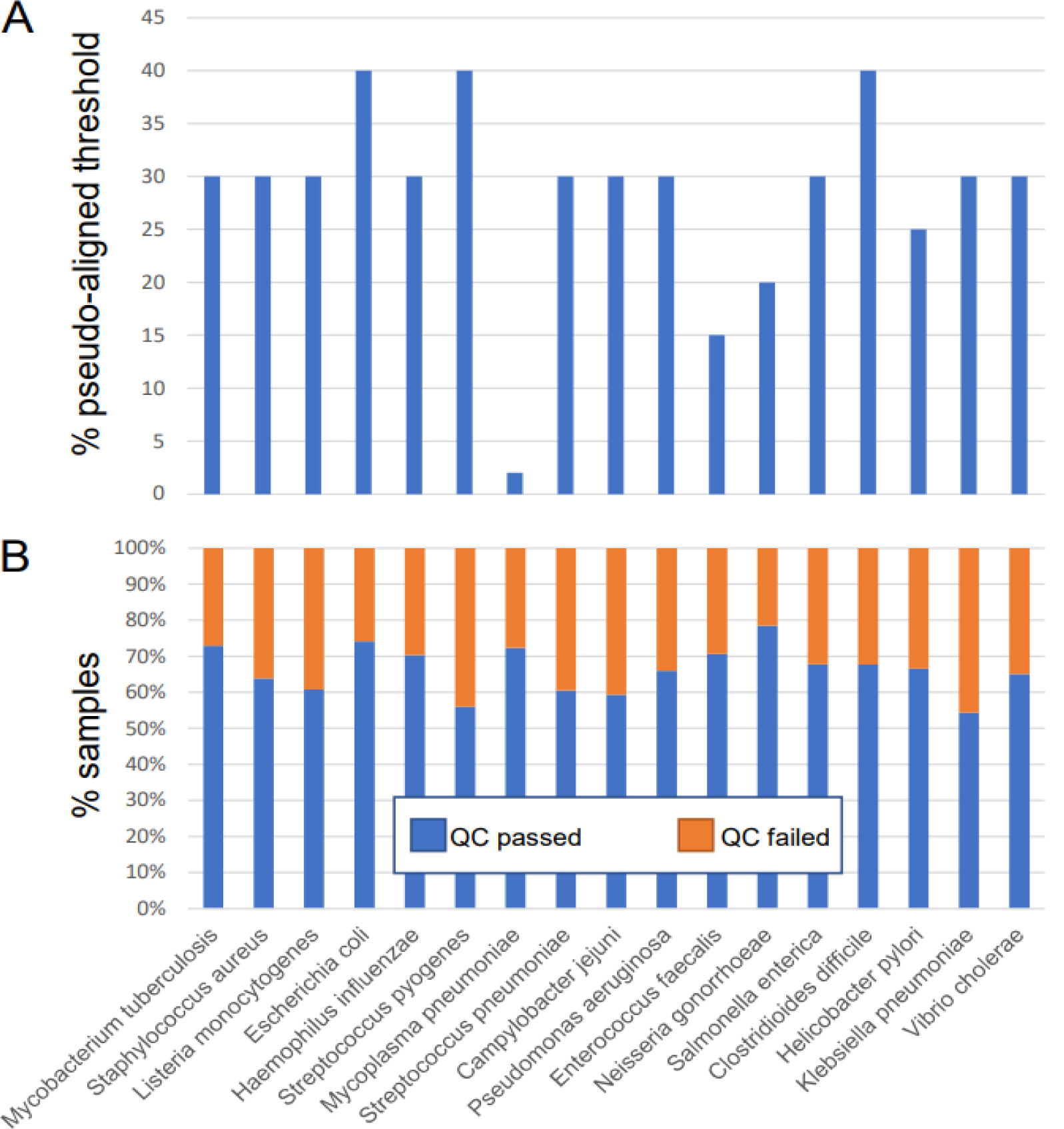
Quality control of the 26270 RNA-seq samples. A) % pseudoaligned (p_pseudoaligned) threshold set for the 17 bacteria. B) The percentage of samples that passed (blue) or failed the p_pseudoaligned an n_pseudoaligned>1000000 thresholds.

### Construction and evaluation of co-expression networks for the 17 bacteria

A small portion of real-world networks is scale-free (Broido and Clauset, 2019), including co-expression networks (Mutwil et al., 2010). In scale-free networks, only a few genes are connected (correlated) to many genes, while the majority of genes show only a few connections (Barabási and Bonabeau, 2003). Scale-free topology is hypothesized to ensure that the network remains mostly unaltered in case of mutations, and is an evolved property that ensures robustness against perturbations (Barabási and Oltvai, 2004). To demonstrate that the expression data of the 17 bacteria can generate biologically meaningful co-expression networks, we investigated whether the data can produce a typical scale-free network. All of the co-expression networks of the 17 bacteria showed a pattern indicative of scale-free topology, as plotting the number of connections a gene has (node degree) against the frequency of this association produced a negative slope (Figure 3A). This confirms the scale-free topology of the co-expression networks and suggests that the networks are biologically relevant.

**Figure 3.**
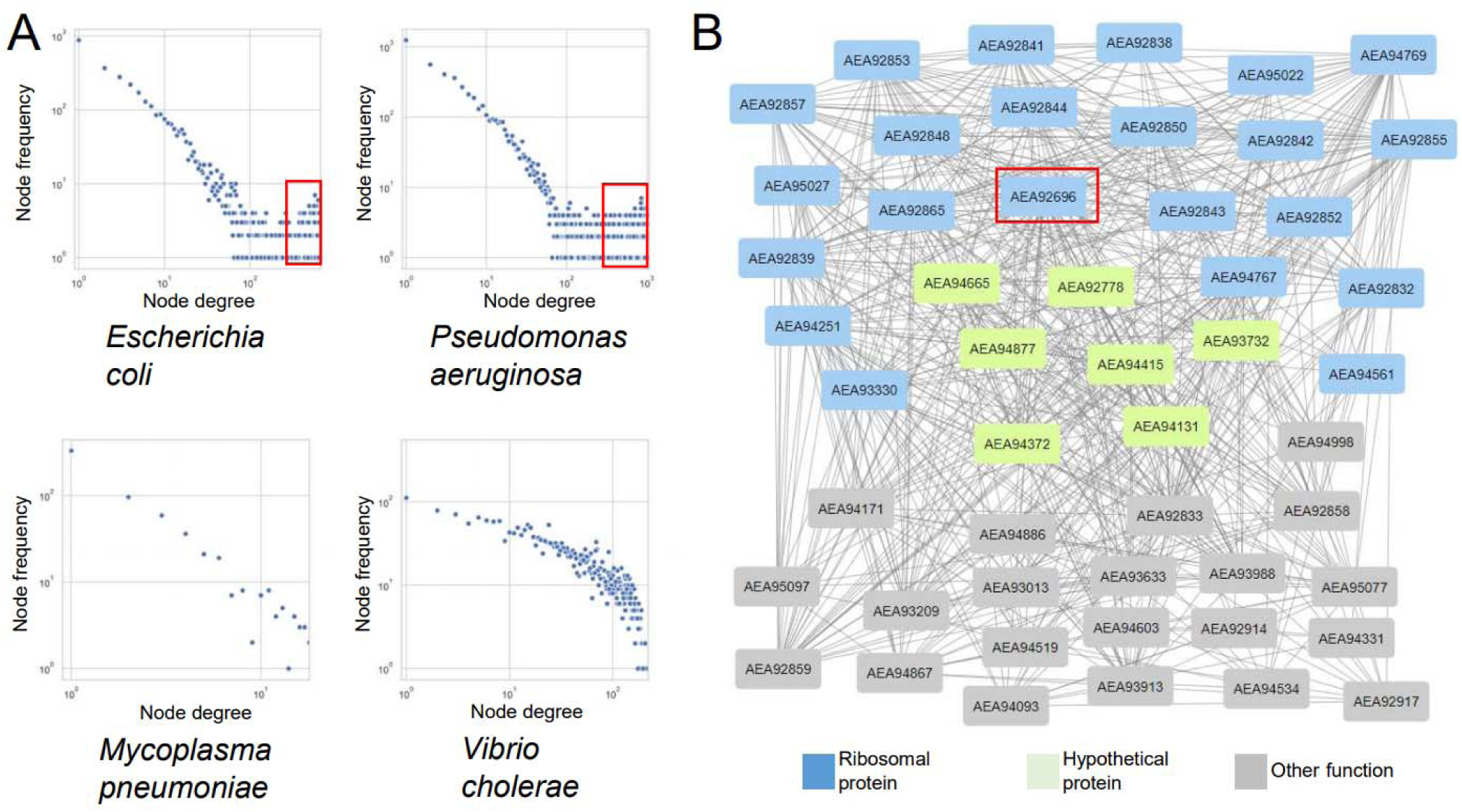
Power-law and example of a ribosomal network. A) Power-law plot obtained from the expression data of the 17 bacteria. The x-axis shows the node degree (number of co-expression connections of a gene), while the y-axis indicates the frequency of a degree. Pearson Correlation Coefficient (PCC) > 0.7 was used to decide whether two genes are co-expressed. The two axes are log10-transformed. B) Co-expression neighborhood of *AEA92696* from *Enterococcus faecalis* (red square), a 30S ribosomal protein S18, and the 50 most highly co-expressed genes (including *AEA92696*). Nodes indicate genes, while gray edges connect genes with PCC > 0.7. Blue nodes represent ribosomal genes, green nodes represent genes with ‘Hypothetical protein’ in their description, while gray nodes indicate genes with other functions.

Interestingly, we observed that the power-law plots of some bacteria contain more nodes with a higher degree than expected from a network following power law (Figure 3A, indicated by red squares). While the basis of this phenomenon is outside of the scope of this publication, we speculate that this is caused by the operon structure of the bacterial genes. Interestingly, certain bacteria, such as *Vibrio cholerae* (Figure 3A) did not show this pattern. Finally, *Mycoplasma pneumoniae* power-law plot showed a small number of points, indicating that few genes show PCC>0.7 in this bacteria. This could be attributed to most samples in this bacterium showing worse mapping statistics than the other 16 bacteria (Figure 2A, Figure S1), indicating that perhaps the available CDS for *Mycoplasma* are of poor quality.

To demonstrate that our co-expression networks can be used to predict novel components of ribosomes, we investigated the co-expression neighborhood of *AEA92696*, a 30S ribosomal protein S18 from *Enterococcus faecalis*. The neighborhood was constructed by retrieving the top 50 genes with the highest PCC values to *AEA92696* (Table S2), where gene pairs with PCC>0.7 are connected (Figure 3B). Out of 50 genes, 22 were annotated as a component of the 30S (e.g., S15, S4, S3) or 50S (e.g., L15, L3, L14) ribosomal subunit, indicating that genes in this neighborhood are involved in protein synthesis. Interestingly, 7 genes in the neighborhood are annotated as ‘hypothetical proteins’ (Figure 3B). Since these genes are found in the neighborhood that is likely to be involved in protein synthesis, we propose that these hypothetical proteins are also involved in protein synthesis in *Enterococcus faecalis*.

### Prediction of novel components of ribosomes by a meta-analysis of the co-expression networks

To predict which genes with unknown function are involved in protein synthesis in the 17 bacteria, we first identified genes that are involved in protein synthesis (search term ‘ribosom’) or shared no similarity to any characterized gene (search term ‘hypothetical’, ‘DUF’, ‘conserved’). The analysis revealed that typically, the ribosomal genes constitute ∼5% of all genes in a bacterial genome (Figure 4A). In comparison, the number of genes that are without functional annotation varies from <1% (*Salmonella enterica*) to 43% (*Helicobacter pylori*).

**Figure 4.**
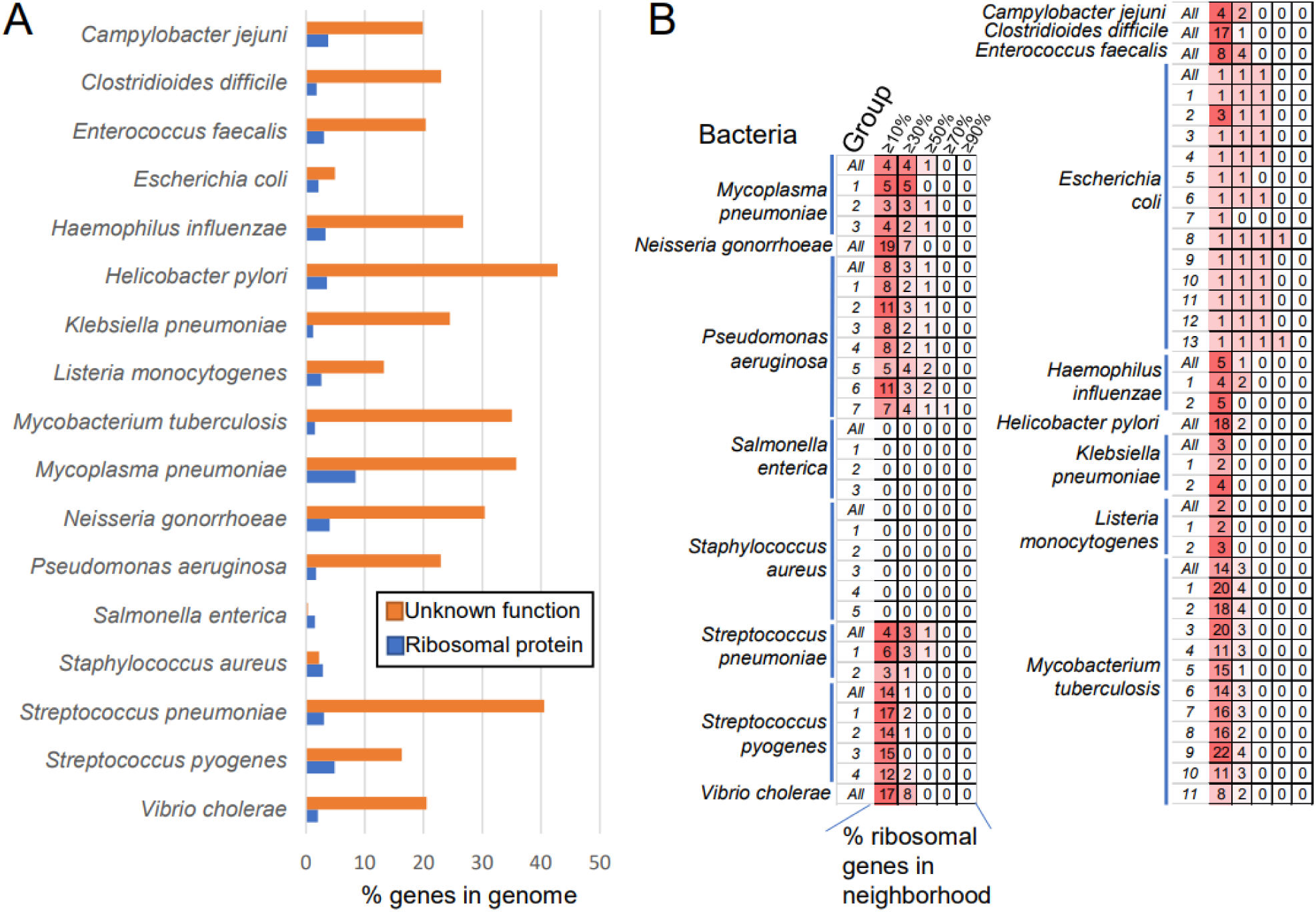
Predicting novel components of ribosomes in the 17 bacteria. A) The percentage of genes wit unknown function (orange bars) and ribosomal genes (blue bars) in the genomes of the 17 bacteria. B) Number of genes with unknown function that are predicted to be involved in protein synthesis in the 17 bacteria. The predictions made by the 60 groups are shown in rows, and the groups are numbered (e.g., *E*.*coli* data is divided into 13 groups). Predictions made on all available data are indicated by ‘All’ in the group column. The last five columns indicate the cutoff that was used to assign a gene with unknown function to protein synthesis. For example, cutoff indicates that at least 10% (5) of genes in the top 50 co-expressed genes were annotated as ribosomal proteins.

Next, to predict uncharacterized genes that are involved in protein synthesis, we calculated the percentage of ribosomal genes that are co-expressed with each uncharacterized gene. More specifically, we looked at the top 50 co-expressed genes of each uncharacterized gene and calculated the percentage of ribosomal genes in this list. The analysis is based on the example shown above (Figure 3B), where a high percentage (43%) of ribosomal genes co-expressed with *AEA92696* indicate that this gene is involved in protein synthesis. We set a percentage threshold that annotates a gene with unknown function as involved in protein synthesis if at least 10% (i.e., at least 5 out of 50 genes), 30%, 50%, 70% or 90% of its top 50 co-expressed genes are involved in protein synthesis. By increasing the threshold, the prediction can be made more stringent, at the cost of the number of genes with the unknown function assigned to protein synthesis (Figure 4B, Table S2).

We observed a varying number of predictions between the different bacteria, ranging from 0 uncharacterized genes assigned to protein synthesis (*Staphylococcus aureus, Salmonella enterica*) to 22 (*Mycobacterium tuberculosis*, Figure 4B). As expected, the number of predictions dropped when the percentage threshold was increased, with almost no genes assigned to protein synthesis at ≥ 50%threshold. Interestingly, we observed a good agreement between the number of predictions made by different student groups. For example, *Mycobacterium tuberculosis* expression data (6495 samples, Table 1) was divided among 11 student groups and used to perform 11 independent predictions (group 1-11), which we compared to a prediction based on the combined data (All). The prediction based on all data (14 uncharacterized genes assigned to protein synthesis at 10% threshold) did not have more (22 predicted genes, group 9) or less (8 predicted genes, group 11). Furthermore, while each group predicted some unique genes, the majority of the predictions identified the same set of genes, indicating that more data is not necessarily better in this scenario.

## Discussion

Protein sequence similarity is commonly used to transfer molecular function annotation from one protein to another (Kulmanov *et al*., 2018). Molecular function annotation by sequence comparison is commonly performed using programs such as BLAST (Altschul *et al*., 1997) and InterProScan (Quevillon *et al*., 2005). However, a substantial proportion of coding sequences lack sequence similarity to any characterized genes (Figure 4)(Rhee and Mutwil, 2014; Ruprecht *et al*., 2017), making sequence similarity-based inference of gene function unsuitable. An excellent example of this limitation are genes that we have analyzed in this study. Since these genes are annotated as ‘hypothetical protein’, ‘domain of unknown function’, or ‘conserved protein’, there is no reference sequence that can be used to elucidate their function.

Transcriptomic data is a rapidly growing resource that captures gene expression levels of all genes in an organism. Co-expression analysis is based on the observation that functionally related genes tend to have similar expression profiles across different experiments, and has become a powerful tool for predicting gene function (Wu *et al*., 2002). We applied this approach to identify novel components of protein synthesis machinery in the 17 most notorious bacteria pathogens, for which sufficient (defined as > 100 RNA-seq samples) expression data exists (Table 1). In this study, we achieved two aims.

Firstly, we show co-expression analysis can be used to predict novel candidates of bacterial ribosomes. We observed that ribosomal proteins tend to be strongly co-expressed (Figure 3B), suggesting that uncharacterized genes co-expressed with the ribosomal proteins are likely involved in some aspect (ribosome assembly, protein synthesis, termination) of protein synthesis. We predicted a substantial number of novel genes involved in protein synthesis for 15 out of 17 bacteria (Figure 4B, Table S3), that can serve as good targets to develop species-specific antibiotics. The available expression data (Supplemental Data 1) for the 17 bacteria can be further mined to study other biological functions and vulnerabilities (e.g., cell wall, RNA, and DNA biosynthesis) of these bacteria.

Secondly, we show that such analysis can be outsourced to a large group of individuals. Here, the gene expression data was streamed and pseudo-aligned by 285 first-year undergraduate students, as part of the Computational Thinking class project. To this end, we used a modified LSTrAP-Cloud pipeline (Tan *et al*., 2020), where the students were divided into 60 groups, and each group was tasked to download and perform quality-control of ∼600 samples over a week (Figure 1-2, Table 1). Theoretically, 85,500 (300*285) RNA-seq samples equivalent to ∼85Tb could be processed per day by the class, providing a computing power rivaling a high-end computer cluster. While each student had access to only two Xeon cores, one of the major bottlenecks in processing the voluminous RNA-seq data, data download, was circumvented by fast internet connection of each Google Colab virtual machine.

While similar approaches are used by, e.g., folding@home, it is to our knowledge the first attempt to process gene expression data in such a manner. We envision that similar approaches will soon allow us to study gene expression data within and across whole kingdoms of life.

## Supplemental figures

**Figure S1. Scatter plot showing the number (x-axis) and percentage (y-axis) of pseudoaligned reads for the 17 bacteria**.

**Figure S2. Power-law plot of the 17 bacteria**. The x-axis shows the node degree (number of coexpression connections of a gene, PCC>0.7), while the y-axis indicates the frequency of a degree. The two axes are log10-transformed.

## Supplemental tables

**Table S1. Quality control of the RNA-seq samples**. The table indicates the species (first column), sample ID (second column), group ID processing the sample (third column), number of pseudoaligned reads (fourth column), percentage of pseudoaligned reads (fifth column) and an indication whether the sample passed the set quality thresholds (sixth column).

**Table S2. Co-expression neighborhood of *AEA92696* from *Enterococcus faecalis***. The genes are sorted according to the Pearson Correlation Coefficient (r, first column). The gene IDs (second column), type (third column, 1 = ribosomal protein, 2 = gene with unknown function, 0 = not 1 or 2) and annotation (fourth column) are indicated.

**Table S3. Uncharacterized genes predicted to be involved in protein synthesis**. Each row contains genes with unknown functions predicted to be involved in protein synthesis. The rows contain predictions made by each group (indicated by numbers) or by all available data (All data). The columns indicate the (i) bacteria, (ii) group ID, (iii-vii) predicted genes at the different percentage thresholds.

## Declarations

The datasets supporting the conclusions of this article are included within the article (and its additional files). The authors declare that they have no competing interests

## Supplemental data

**Supplemental Data 1. Expression matrices for 17 bacteria.**

## Acknowledgments

We would like to thank Google for providing Google Colab, and all members of the class of BS1009 (Introduction to Computational Thinking) that ran during the 2019/20 academic year.

## References

Ahmed, T., Shi, J. and Bhushan, S. (2017) Unique localization of the plastid-specific ribosomal proteins in the chloroplast ribosome small subunit provides mechanistic insights into the chloroplastic translation. Nucleic Acids Res.

Ahmed, T., Yin, Z. and Bhushan, S. (2016) Cryo-EM structure of the large subunit of the spinach chloroplast ribosome. Sci. Rep.

Altschul, S.F., Madden, T.L., Schäffer, A.A., Zhang, J., Zhang, Z., Miller, W. and Lipman, D.J. (1997) Gapped BLAST and PSI-BLAST: A new generation of protein database search programs. Nucleic Acids Res.

Arenz, S. and Wilson, D.N. (2016) Bacterial protein synthesis as a target for antibiotic inhibition. Cold Spring Harb. Perspect. Med.

Barabási, A.-L. and Bonabeau, E. (2003) Scale-free networks. Sci. Am., 288, 60–9.

Barabási, A.L. and Oltvai, Z.N. (2004) Network biology: Understanding the cell’s functional organization. Nat. Rev. Genet., 5, 101–113.

Barandun, J., Hunziker, M., Vossbrinck, C.R. and Klinge, S. (2019) Evolutionary compaction and adaptation visualized by the structure of the dormant microsporidian ribosome. Nat. Microbiol.

Bieri, P., Leibundgut, M., Saurer, M., Boehringer, D. and Ban, N. (2017) The complete structure of the chloroplast 70S ribosome in complex with translation factor pY. EMBO J.

Bray, N.L., Pimentel, H., Melsted, P. and Pachter, L. (2016) Near-optimal probabilistic RNA-seq quantification. Nat. Biotechnol., 34, 525–527.

Broido, A.D. and Clauset, A. (2019) Scale-free networks are rare. Nat. Commun.

Dinos, G.P. (2017) The macrolide antibiotic renaissance. Br. J. Pharmacol.

Dunkle, J.A., Xiong, L., Mankin, A.S. and Cate, J.H.D. (2010) Structures of the Escherichia coli ribosome with antibiotics bound near the peptidyl transferase center explain spectra of drug action. Proc. Natl. Acad. Sci. U. S. A.

Eyal, Z., Matzov, D., Krupkin, M., Wekselman, I., Paukner, S., Zimmerman, E., Rozenberg, H., Bashan, A. and Yonath, A. (2015) Structural insights into species-specific features of the ribosome from the pathogen Staphylococcus aureus. Proc. Natl. Acad. Sci. U. S. A.

Ferrari, C., Proost, S., Ruprecht, C. and Mutwil, M. (2018) PhytoNet: Comparative co-expression network analyses across phytoplankton and land plants. Nucleic Acids Res., 46, W76–W83.

Golkar, T., Zielinski, M. and Berghuis, A.M. (2018) Look and outlook on enzyme-mediated macrolide resistance. Front. Microbiol.

Greber, B.J. and Ban, N. (2016) Structure and Function of the Mitochondrial Ribosome. Annu. Rev. Biochem.

Hansen, B.O., Meyer, E.H., Ferrari, C., Vaid, N., Movahedi, S., Vandepoele, K., Nikoloski, Z. and Mutwil, M. (2018) Ensemble gene function prediction database reveals genes important for complex I formation in Arabidopsis thaliana. New Phytol., 217, 1521–1534.

Hansen, B.O., Vaid, N., Musialak-Lange, M., Janowski, M. and Mutwil, M. (2014) Elucidating gene function and function evolution through comparison of co-expression networks of plants. Front. Plant Sci., 5.

Jiménez-Gómez, J.M., Wallace, A.D. and Maloof, J.N. (2010) Network analysis identifies ELF3 as a QTL for the shade avoidance response in arabidopsis. PLoS Genet., 6.

Kerr, I.D., Reynolds, E.D. and Cove, J.H. (2005) ABC proteins and antibiotic drug resistance: Is it all about transport? Biochem. Soc. Trans.

Kulmanov, M., Khan, M.A. and Hoehndorf, R. (2018) DeepGO: Predicting protein functions from sequence and interactions using a deep ontology-aware classifier. Bioinformatics, 34, 660–668.

Kushwaha, A.K. and Bhushan, S. (2020) Unique structural features of the Mycobacterium ribosome. Prog. Biophys. Mol. Biol.

Lee, I., Ambaru, B., Thakkar, P., Marcotte, E.M. and Rhee, S.Y. (2010) Rational association of genes with traits using a genome-scale gene network for Arabidopsis thaliana. Nat. Biotechnol., 28, 149–156. Available at: http://dx.doi.org/10.1038/nbt.1603.

Leinonen, R., Sugawara, H. and Shumway, M. (2011) The sequence read archive. Nucleic Acids Res., 39.

Lin, J., Zhou, D., Steitz, T.A., Polikanov, Y.S. and Gagnon, M.G. (2018) Ribosome-Targeting Antibiotics: Modes of Action, Mechanisms of Resistance, and Implications for Drug Design. Annu. Rev. Biochem.

Liu, M. and Douthwaite, S. (2002) Activity of the ketolide telithromycin is refractory to Erm monomethylation of bacterial rRNA. Antimicrob. Agents Chemother.

Mah, T.F.C. and O’Toole, G.A. (2001) Mechanisms of biofilm resistance to antimicrobial agents. Trends Microbiol., 9, 34–39.

Melnikov, S., Ben-Shem, A., Garreau De Loubresse, N., Jenner, L., Yusupova, G. and Yusupov, M. (2012) One core, two shells: Bacterial and eukaryotic ribosomes. Nat. Struct. Mol. Biol.

Melnikov, S., Manakongtreecheep, K. and Söll, D. (2018) Revising the structural diversity of ribosomal proteins across the three domains of life. Mol. Biol. Evol.

Murina, V., Kasari, M., Takada, H., et al. (2019) ABCF ATPases Involved in Protein Synthesis, Ribosome Assembly and Antibiotic Resistance: Structural and Functional Diversification across the Tree of Life. J. Mol. Biol.

Mutwil, M., Klie, S., Tohge, T., et al. (2011) PlaNet: Combined sequence and expression comparisons across plant networks derived from seven species. Plant Cell, 23, 895–910. Available at: http://www.plantcell.org/cgi/doi/10.1105/tpc.111.083667.

Mutwil, M., Obro, J., Willats, W.G.T. and Persson, S. (2008) GeneCAT--novel webtools that combine BLAST and co-expression analyses. Nucleic Acids Res., 36.

Mutwil, M., Ruprecht, C., Giorgi, F.M., Bringmann, M., Usadel, B. and Persson, S. (2009) Transcriptional wiring of cell wall-related genes in Arabidopsis. Mol. Plant, 2, 1015–1024.

Mutwil, M., Usadel, B., Schütte, M., Loraine, A., Ebenhöh, O. and Persson, S. (2010) Assembly of an interactive correlation network for the Arabidopsis genome using a novel Heuristic Clustering Algorithm. Plant Physiol., 152, 29–43.

Ng, J.W.X., Tan, Q.W., Ferrari, C. and Mutwil, M. (2019) Diurnal.plant.tools: Comparative Transcriptomic and Co-expression Analyses of Diurnal Gene Expression of the Archaeplastida Kingdom. Plant Cell Physiol.

Patro, R., Duggal, G., Love, M.I., Irizarry, R.A. and Kingsford, C. (2017) Salmon provides fast and bias-aware quantification of transcript expression. Nat. Methods, 14, 417–419.

Proost, S. and Mutwil, M. (2018a) CoNekT: An open-source framework for comparative genomic and transcriptomic network analyses. Nucleic Acids Res., 46, W133–W140. Available at: https://xswww.biorxiv.org/content/early/2018/01/28/255075?rss=1.

Proost, S. and Mutwil, M. (2018b) CoNekT: An open-source framework for comparative genomic and transcriptomic network analyses. Nucleic Acids Res., 46, W133–W140. Available at: https://academic.oup.com/nar/article/46/W1/W133/4990637.

Proost, S. and Mutwil, M. (2017) Planet: Comparative co-expression network analyses for plants. In A. D. J. van Dijk, ed. Methods in Molecular Biology. New York, NY: Springer New York, pp. 213–227. Available at: http://dx.doi.org/10.1007/978-1-4939-6658-5_12.

Proost, S. and Mutwil, M. (2016) Tools of the trade: Studying molecular networks in plants. Curr. Opin. Plant Biol., 30, 130–140.

Quevillon, E., Silventoinen, V., Pillai, S., Harte, N., Mulder, N., Apweiler, R. and Lopez, R. (2005) InterProScan: Protein domains identifier. Nucleic Acids Res., 33.

Rhee, S.Y. and Mutwil, M. (2014) Towards revealing the functions of all genes in plants. Trends Plant Sci., 19, 212–221.

Ruprecht, C., Mendrinna, A., Tohge, T., Sampathkumar, A., Klie, S., Fernie, A.R., Nikoloski, Z., Persson, S. and Mutwil, M. (2016) Famnet: A framework to identify multiplied modules driving pathway expansion in plants. Plant Physiol., 170, 1878–1894. Available at: http://www.plantphysiol.org/lookup/doi/10.1104/pp.15.01281.

Ruprecht, C., Mutwil, M., Saxe, F., Eder, M., Nikoloski, Z. and Persson, S. (2011) Large-scale co-expression approach to dissect secondary cell wall formation across plant species. Front. Plant Sci., 2, 1–13.

Ruprecht, C., Proost, S., Hernandez-Coronado, M., Ortiz-Ramirez, C., Lang, D., Rensing, S.A., Becker, J.D., Vandepoele, K. and Mutwil, M. (2017) Phylogenomic analysis of gene co-expression networks reveals the evolution of functional modules. Plant J., 90, 447–465.

Sharkey, L.K.R. and O’Neill, A.J. (2018) Antibiotic Resistance ABC-F Proteins: Bringing Target Protection into the Limelight. ACS Infect. Dis.

Sibout, R., Proost, S., Hansen, B.O., et al. (2017) Expression atlas and comparative coexpression network analyses reveal important genes involved in the formation of lignified cell wall in Brachypodium distachyon. New Phytol., 215, 1009–1025.

Spellberg, B., Blaser, M., Guidos, R.J., et al. (2011) Combating antimicrobial resistance: Policy recommendations to save lives. Clin. Infect. Dis., 52.

Stuart, J.M., Segal, E., Koller, D. and Kim, S.K. (2003) A gene-coexpression network for global discovery of conserved genetic modules. Science (80-.)., 302, 249–255. Available at: http://www.ncbi.nlm.nih.gov/pubmed/12934013.

Takabayashi, A., Ishikawa, N., Obayashi, T., Ishida, S., Obokata, J., Endo, T. and Sato, F. (2009) Three novel subunits of Arabidopsis chloroplastic NAD(P)H dehydrogenase identified by bioinformatic and reverse genetic approaches. Plant J., 57, 207–219.

Takahashi, N., Lammens, T., Boudolf, V., Maes, S., Yoshizumi, T., Jaeger, G. De, Witters, E., Inzé, D. and Veylder, L. De (2008) The DNA replication checkpoint aids survival of plants deficient in the novel replisome factor ETG1. EMBO J., 27, 1840–1851. Available at: http://www.ncbi.nlm.nih.gov/pubmed/18528439 [Accessed February 17, 2017].

Tan, Q.W., Goh, W. and Mutwil, M. (2020) LSTrAP-Cloud: A User-friendly Cloud Computing Pipeline to Infer Co-functional and Regulatory Networks. bioRxiv.

Tan, Q.W. and Mutwil, M. (2019) Inferring biosynthetic and gene regulatory networks from Artemisia annua RNA sequencing data on a credit card-sized ARM computer. Biochim. Biophys. acta. Gene Regul. Mech., 194429.

Usadel, B., Obayashi, T., Mutwil, M., et al. (2009) Co-expression tools for plant biology: Opportunities for hypothesis generation and caveats. Plant, Cell Environ., 32, 1633–1651.

Vazquez-Laslop, N., Thum, C. and Mankin, A.S. (2008) Molecular Mechanism of Drug-Dependent Ribosome Stalling. Mol. Cell.

Wen Tan, Q. and Mutwil, M. (2019) Malaria.tools—comparative genomic and transcriptomic database for Plasmodium species. Nucleic Acids Res.

Wilson, D.N. (2014) Ribosome-targeting antibiotics and mechanisms of bacterial resistance. Nat. Rev. Microbiol.

Wilson, D.N. (2009) The A-Z of bacterial translation inhibitors. Crit. Rev. Biochem. Mol. Biol.

Wu, L.F., Hughes, T.R., Davierwala, A.P., Robinson, M.D., Stoughton, R. and Altschuler, S.J. (2002) Large-scale prediction of Saccharomyces cerevisiae gene function using overlapping transcriptional clusters. Nat. Genet., 31, 255–265. Available at: http://www.nature.com/doifinder/10.1038/ng906.

Yu, H., Luscombe, N.M., Qian, J. and Gerstein, M. (2003) Genomic analysis of gene expression relationships in transcriptional regulatory networks. Trends Genet., 19, 422–427.

