## Supplementary figures and images for "LSTrAP-Crowd: Prediction of novel components of bacterial ribosomes with crowd-sourced analysis of RNA sequencing data"

### Figure S1

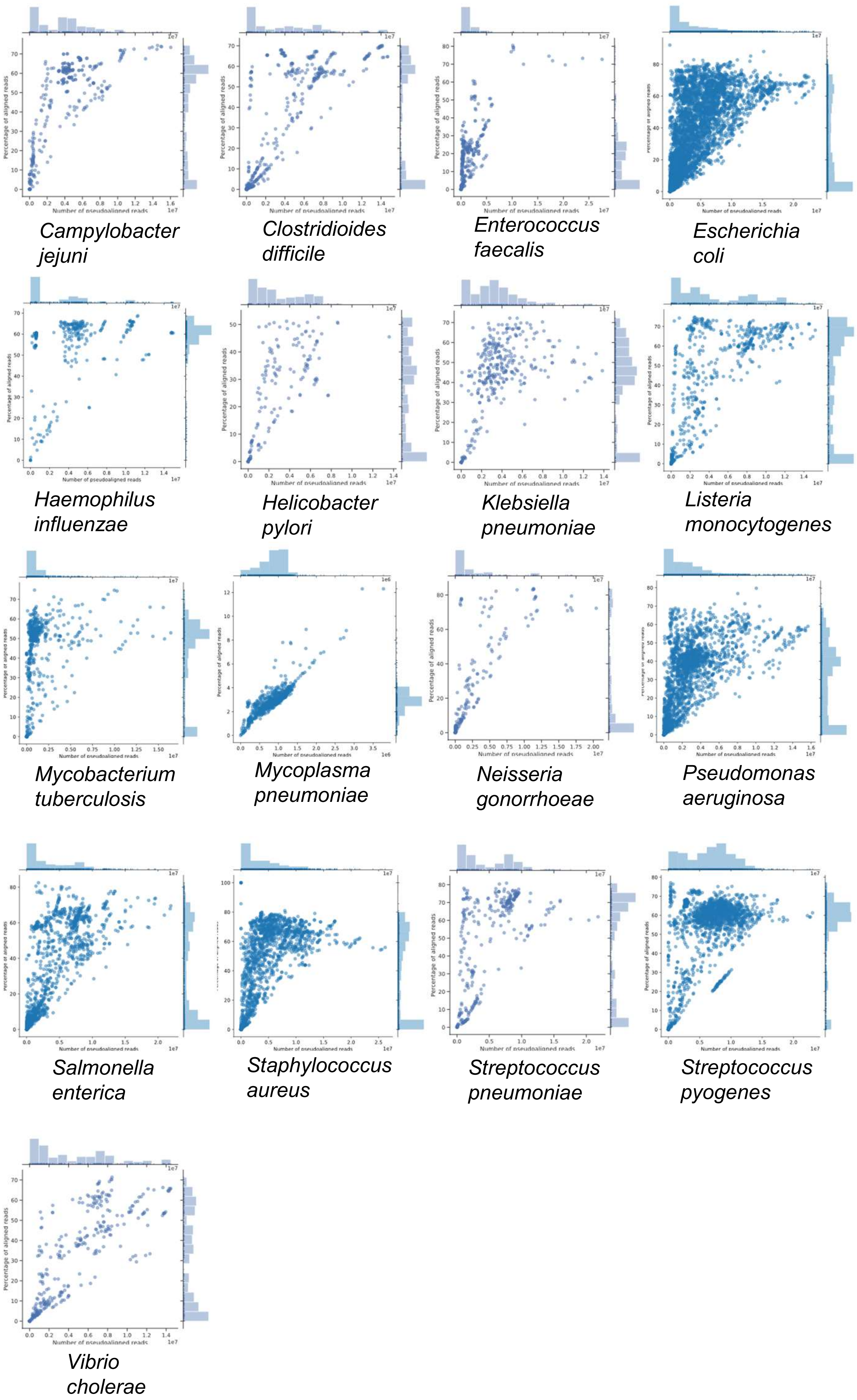

### Figure S2

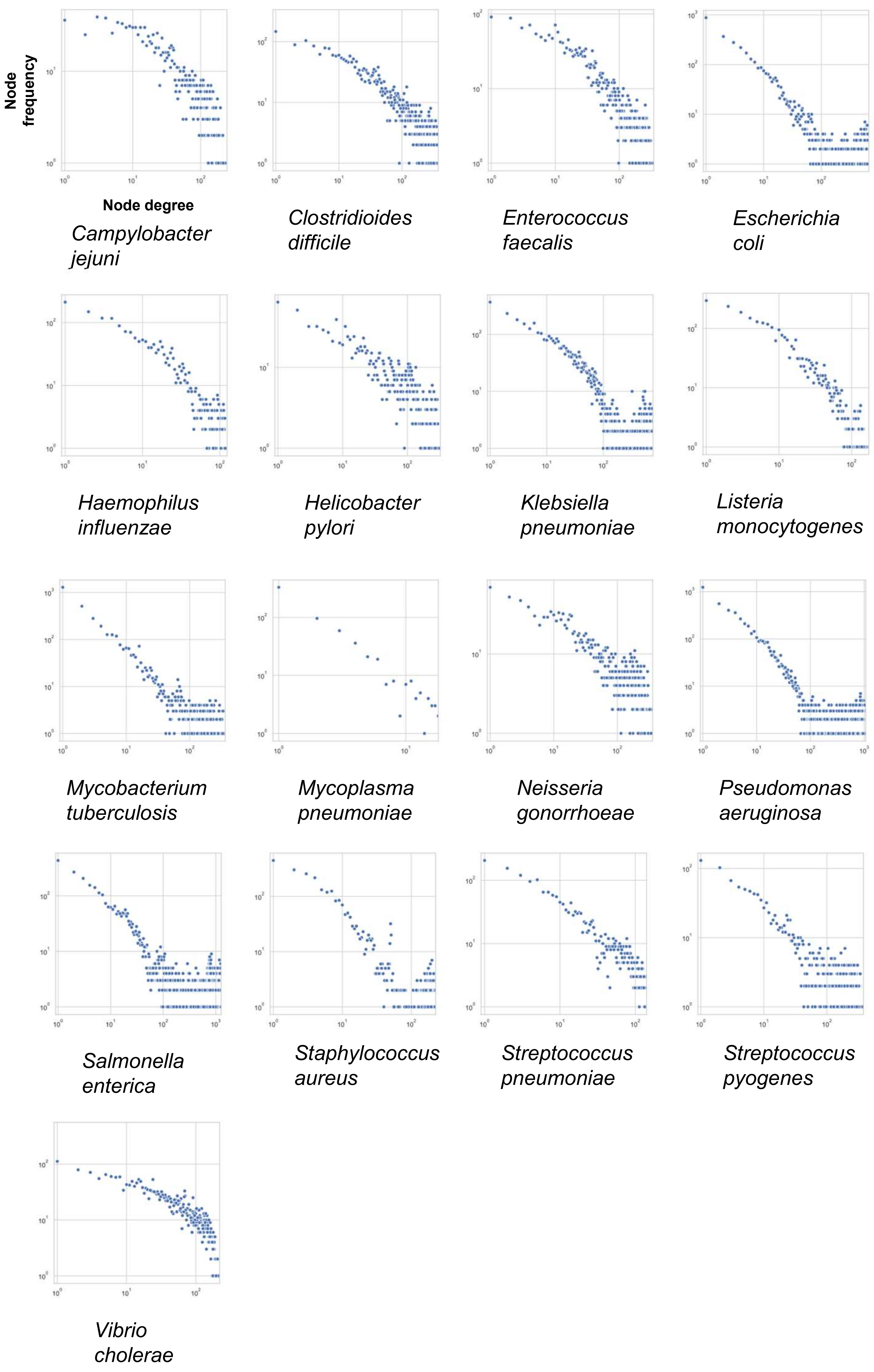
